# An ability to respond begins with inner alignment: How phase synchronisation effects transitions to higher levels of agency

**DOI:** 10.1101/2024.02.16.580248

**Authors:** Tazzio Tissot, Mike Levin, Chris Buckley, Richard Watson

## Abstract

How do multiple active components at one level of organisation create agential wholes at higher levels of organisation? For example, in organismic development, how does the multi-scale autonomy of the organism arise from the interactions of the molecules, cells and tissues that an organism contains? And, in the major evolutionary transitions, how does a multicellular organism, for example, arise as an evolutionary unit from the selective interests of its unicellular ancestors? We utilise computational models as a way to think about this general question. We take a deliberately minimalistic notion of an agent: a competency to take one of two possible actions to minimise stress. Helping ourselves to this behaviour at the microscale, we focus on conditions where this same type of agency appears spontaneously at a higher level of organisation. We find that a simple process of positive feedback on the timing of individual responses, loosely analogous to the natural phase synchronisation of weakly coupled oscillators, causes such a transition in behaviour. The emergent collectives that arise become, quite suddenly, able to respond to their external stresses in the same (minimal) sense as the original microscale units. This effects a dramatic rescaling of the system behaviour, and a quantifiable increase in problem-solving competency, serving as a model of how higher-level agency emerges from a pool of lower-level agents or active matter. We discuss how this dynamical ‘waking-up’ of higher-level collectives, through the alignment of their internal dynamics, might relate to reproductive/cell-cycle synchronisation in evolutionary transitions and development.

## Introduction

How do many become one (1-5)? At many different levels of organisation from molecular biology to human society, multiple individuals can sometimes change from acting separately, with individual goals and competencies, to working together in a way that produces a new entity, behaving as a single agent at a higher level of organisation, with new goals and new competencies (1, 2, 6-8). Each of us, for example, is a collection of cells (each retaining a repertoire of semi-autonomous behaviours and functions), that work together to create and maintain a coherent whole capable of collective action and goal-directed behaviour (2). In the developmental emergence of an individual from the active matter of which it is composed, the salient level of agency repeatedly transitions from one level of organisation to another, pivoting through different dynamic and behavioural spaces, including gene-expression, metabolic activity, cellular spatial coordination, electrical excitation (of neural and aneural cells), to large-scale behavioural alignment and coordinated action at the organismic level (1). At a quite different timescale, the process of evolution by natural selection has also scaled-up through multiple levels of organisation; rescaling the ability to exhibit heritable variation in reproductive success from parts to wholes recursively (9-11). These include transitions from multiple self-replicating molecules to the first chromosomes, the first protocells with multiple chromosomes, eukaryote cells with multiple previously free-living organelles, to multicellular organisms, to eusocial colonies (12-15).

These transitions in organismal and evolutionary agency are intimately related. Creation of a higher-level evolutionary unit might partly explain why components work together so well in organismic development and behaviour. On the other hand, the causal relationship may well be the other way around; i.e. it is the change in the level of organismal agency, when directed at the competency of survival and reproduction, which originates the new level of evolutionary unit on which selection can act (10). Understanding how new levels of agency arise (including new levels of organismic competency in developmental time and transitions in the evolutionary unit over evolutionary time) is an open problem (1, 2, 9, 16-18).

Agency shares some conceptual problems with non-reductionist accounts more generally.

Although any behaviour of an agent can be traced to the behaviours of its component parts, it is also quite obvious that the same parts may behave differently when differently organised. Whilst the apparent generality and completeness of lower-level explanations can be attractive, ignoring the circumstances by which higher-level organisations constrain the ‘options’ or offer new ‘affordances’ to the component parts tends to make-invisible the biology that is interesting (1, 7, 9, 18, 19).

We wish to understand, in a simple mechanistic way, how multiple individuals can co-create the necessary organisational conditions to cause them to work together; in particular, when doing so appears to oppose the self-interest of component members. This transitional way of thinking about agency turns focus away from trying to define a categorical notion of agency, to draw a line in the sand, sufficient to distinguish between non-agential and agential systems (18). Instead, it focusses on how a given competency at one level of organisation transitions into a (perhaps similar) competency at another higher level of organisation (9, 10, 20). This can still be an interesting question even when the notion of competency employed is relatively simple. In particular, we consider the competency to take an action (from a finite set) that optimises some quantity, e.g. maximises utility (or reward or fitness), or minimises stress (or frustration or energy), in response to a small signal in this quantity. A mechanism for this notion of agency is not difficult to satisfy at the level of one unit; even a very simple physical model such as a ball rolling downhill over a landscape (with a suitably shaped surface, as we will detail) is sufficient. But the interesting question is whether a collection of units with this competency can organise themselves to create a higher-level agent with comparable competency (21-23). How do multiple self-interested agents at one level of organisation, work together to create agency at a higher level of organisation, thus serving the long-term collective interest of the whole, even when this over-rules the short-term self-interest of the parts? (20).

We thus seek a microscale mechanistic model, complete in determining the behaviour of the system. But we seek to understand a phenomenon that is interesting precisely because a macroscale mechanistic process emerges that changes the outcomes of the system. What does it mean to be a macroscale process if the microscale process completely defines the dynamics of the model? Is it not just an explanatorily-redundant epiphenomenon of the microscale process? These questions are at the heart of agency debates (18). They arise in organismal development and what it means to be a ‘self’ (2), and in evolutionary transitions and what it means to be an evolutionary unit (9, 10, 13, 24). In this paper, we build a very simple and abstract model to investigate these questions. Whilst the example model we present omits many features desirable in a general model of agency, it enables us to address some of these conceptual questions clearly.

Our methodology is to use computational models, not as accounts of, or substitutes for, empirical study, but as tools to think with – to explore the logical consequences of different abstract scenarios and general assumptions (25). What follows therefore is not a specific model of development, nor of evolutionary transitions. Rather it is a model that illustrates some general underlying principles involved in multi-scale competencies and transitions, to help us think about what that might mean, and how to make sense of some conceptual difficulties.

We use this approach to explore the hypothesis that a transition in agency can occur through adjustments in the *timing* of decisions. We model a number of agents who each take actions in a ‘decision cycle’ that repeatedly changes between undecided and decided states. We show that when these decision cycles are not appropriately organised, a collection of agents is unable to exhibit collective action. We illustrate this in a simple scenario where the behaviour of each agent is constrained by pairwise interactions with other agents. Through individual actions, the system quickly reaches a local Nash equilibrium where, even though there exist configurations where many more interactions could be simultaneously satisfied, any unilateral change in the action of a single agent is opposed by interactions with others. Changing from one such equilibrium to another by individual action is therefore not possible (by definition of a local equilibrium) and changing by stochastic multi-agent variations is exponentially unlikely in the distance between equilibria (i.e. requires many specific and simultaneous state changes). In contrast, here we show that when the decision cycles of particular agents are synchronised and the members of this collective enter the undecided state at the same time, the collective exhibits, in this moment, an increased sensitivity to the environment outside the collective, and is thereby able to respond to influences external to the collective and exhibit mass-action that was otherwise unavailable – i.e. *an ability to respond begins with inner alignment*. This changes the configuration space of behaviours (17), rescaling the dynamics of the system from individual actions to specific collective actions that enable the system to move (in a deterministic, not stochastic, fashion) from one equilibrium to another, moving toward states that resolve more interactions. The structure of this synchronisation (i.e. who is synchronised with whom) implicitly defines the membership of the collective and the identity of the higher-level agent. This would not be a very interesting result if we assumed foreknowledge of who belongs together in a group – the membership of each collective. But importantly, we do not assume this but show that the necessary synchronisation structure can be produced using only individual-based, local information, ‘bottom-up’. Changes in timing do not use foreknowledge of, or any sampling of, the benefit attained by the collective action enabled when the group is fully synchronised. Nonetheless, the collective action that results is specific in enabling the system to change from one equilibrium to another, without visiting intermediate states that are opposed by individual action. In this sense, the model demonstrates a bottom-up transition in agency, exhibiting a new competency at the higher-level without presupposing higher-level selection or a higher-level utility-maximisation process, and demonstrating collective actions.

In the following section we clarify the ‘transitions problem’ used to illustrate this effect and describe the model of agents and their decision cycles. We then demonstrate the ability to produce a transition in agency, and the change in problem-solving competency that this confers. We provide two versions of the model: one which models utility-maximising strategic agents and one which models fitness-maximising populations. In both cases, the synchronisation that occurs between the decision-cycles of low-level agents under utility/fitness maximisation behaves much like the spontaneous phase synchronisation that occurs between weakly-coupled oscillators (26). We then discuss some implications of our findings for organismic development and its interactions with evolutionary transitions in individuality. Finally, we use this model to discuss some of the conceptual problems of micro- and macro-scale mechanistic processes, and their interpretation for transitions in agency.

## Model

### The transitions problem: Conflict between levels of agency

To concretise the issues involved, we utilise a simple modular coordination game. A familiar example is the driving conventions problem (Box 1). It is easy for the drivers within each country to arrive at a consensus by each driving on the side that minimises the collisions they experience (hence, agreeing with the majority of others in that country). But, as is familiar in the real world, it may be the case that different countries arrive at a different consensus. At this locally optimal state, each driver suffers no collisions with others in their own country, but suffers occasional collisions with foreigners.

#### Box 1.

**The driving conventions problem**

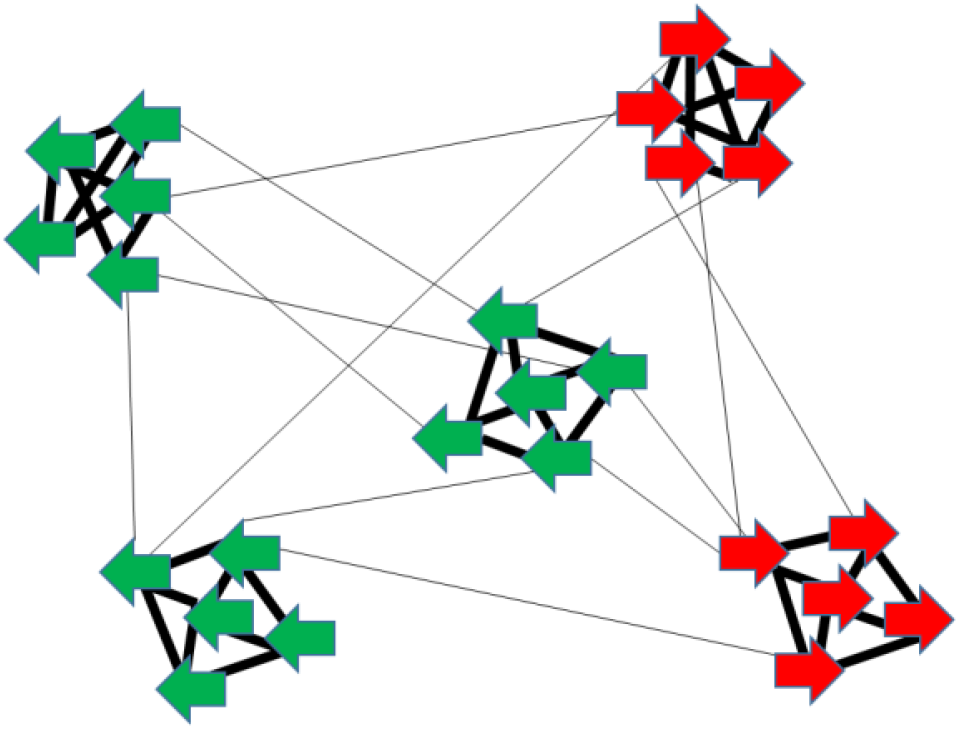

Drivers in the same cluster (country) meet one another much more often (thick lines), than drivers in different countries (thin lines) (not all interactions shown). Each individual driver has the competency to drive on the side that minimises the collisions/stress they experience. This is sufficient to produce a *local* consensus (either all left/green or all right/red depending on the local majority). However, this is only a locally optimal state for the system as a whole since all drivers experience some (less frequent) collisions with drivers in other countries. No one driver can change their decision unilaterally to reduce these collisions. Even a driver in the global minority (red drivers in this example) cannot change because, if acting alone, the cost of increasing their collisions with locals is greater than the benefit of decreasing collisions with foreigners. Two global optima exist, where all collisions are avoided. To escape a local optimum and arrive at (or move toward) a global consensus requires individuals to move against their short-term self-interest (increasing stress/collisions). Note that, if only all the drivers within a country were capable of acting together such that they all changed simultaneously in the right direction (toward the global majority) this would satisfy more external conflicts without creating any internal conflicts. But although an individual driver has the competency to respond to the influence of other drivers, a country is not an agent that has this competency. That is, although a modular constraint structure is part of the problem definition, the agents do not know this structure or who they need to act together with, and a country is not a level of organisation that is able to respond to the influence of its environment (including other countries). It is locked in place, trapped at the local equilibrium, by the pull of its internal stresses. (See Discussion for biological interpretations).

Just like individuals within a country, countries are (in the sense of an average over their components) incentivised to agree with the majority of other countries to reduce collective stress. But just because a country is composed of agential members does not mean that the country is also agential. A country does not have the same agential competency that an individual driver has (minimal though that is); a country is not a thing that is able to take decisions or actions. Intuitively, after arriving at a local equilibrium, a country is unable to ‘change its mind’ and move to a different one. How can such collectives acquire the ability to take collective actions that minimise collective stress (precisely those that are composed of many individual actions that, if taken unilaterally, would *increase* stress)?

For a modular driving conventions problem the interaction strengths (or constraints between drivers) can be defined as follows:

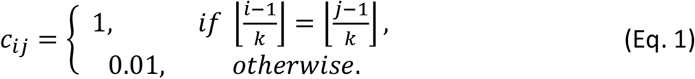

where *i* and *j*∈{1, … *N*} are the indexes of the pairwise constraints over *N* variables, ⌊*x*⌋ is the floor of the value x, and *k* is the number of variables in each module (‘country’).

In the following experiments, we use *N*=300 and *k*=10, creating 30 countries of 10 drivers each. Following Hopfield (27, 28), it is useful for such problems to define an energy function describing the extent to which all the constraints in the system are satisfied by the different combinations of discretised unit states:

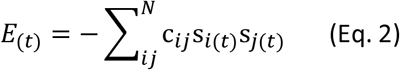

When the interaction terms are symmetric (*c*_*ij*_ = *c*_*ji*_), as per Eq. 1, the behaviour of a system with discrete states is described by state changes that effect the local minimisation of this energy function, and is guaranteed to arrive at a fixed point (27). Such points correspond to a Nash equilibrium where no unit can increase their utility or reduce stress by changing state unilaterally^1^. This means that all drivers easily find agreement with the other drivers within their own country, but not necessarily with those in other countries. With these parameters, there are 2^30^ different local optima (i.e. the drivers in each of 30 countries will reach one of 2 conventions). Only two of these is globally optimal (i.e. when all drivers in all countries find agreement) (Fig. 1).

**Figure 1.**
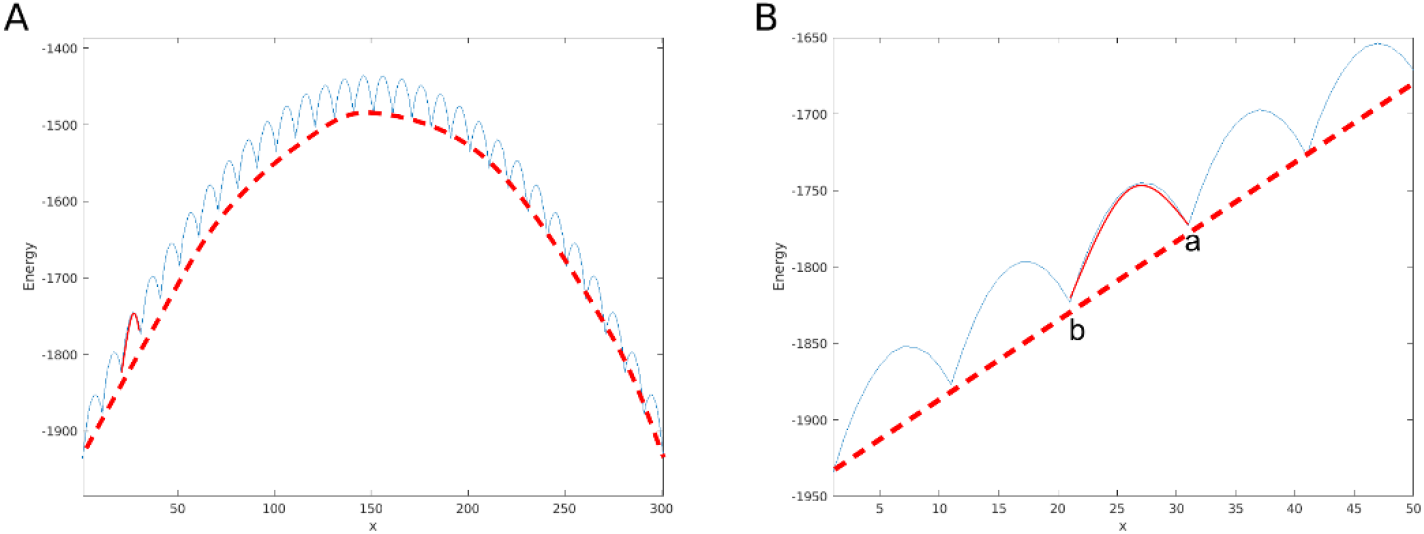
The energy (or stress) landscape of the driving convention problem. (A) A cross-section (blue) through the energy surface (Eq. 2) from one global minimum (all Left) to the other (all Right) (through the state configurations described by the regular expression L^(N-x)^R^x^) shows some of the 2^30^ locally optimal configurations. The red dotted curve indicates the energy contribution from only the (weaker) between-country interactions. The solid red curve is an example of the energy contribution from only the within-country interactions. The complete energy surface is a sum of these local and global components. (B) A close-up of part of the energy surface. Notice that between point **a** (one of the local optima) and point **b** (the closest local optimum to **a** in the direction of the global optimum) there are 10 units that need to change state. To change one unit at a time would mean climbing up the energy barrier caused by the strong within-country interactions (solid red), i.e. this moves against short-term self-interest (increasing stress) even though it would serve their long-term collective interest to be at **b**. Notice that if somehow this energy barrier were temporarily removed there would remain a downward gradient between **a** and **b** caused by the between-country interactions.

Considering this kind of two-scale energy function^2^, one way to understand our aim is to describe a dynamical system that changes how it behaves from one scale to the other: before the transition it follows individual gradients (and is unable to follow the global gradient), whereas after the transition it follows the global gradient (ignoring individual gradients). A system that transitions in this way must exhibit specific coordinated collective action to overcome the pull of the individual-level stress gradient, and in so doing will be able to move through the energy landscape to the global optimum unhindered. This gives an outcome equivalent to actions that favour long-term collective outcomes even though these over-rule the short-term individual outcomes that otherwise trap the system at a local equilibrium.

The possibility of long-term collective gains suggests that if only some suitably structured relationships could be put in place, to alter the short-term incentives or responses of the parts in the right way, these would be to everyone’s benefit in the long-term. For example, mandated driving regulations (that over-rule individual choices). However, such relationships need to coordinate actions among many parts simultaneously. A partially constructed relationship structure can therefore be worse than none because changing only some of the parts (e.g. a country with a fraction of the drivers using the foreign convention) is worse than staying as you are. So even if such relationships or incentive modifiers (see also ‘individuating traits’ in an evolutionary process (29)) were identifiable in principle, and beneficial when fully established, this does not on its own explain how self-interested individuals could create such organised relationships bottom-up through individual incentives (10). A bottom-up account is necessary because a microscale mechanistic model for transitions in agency must operate without presupposing the higher-level agency it intends to explain.

Note that the existence of locally optimal states depends on the relative frequency of intra-cluster and inter-cluster interactions. If the rate of inter-cluster interactions is assumed to be sufficiently high, then there are no local optima in the dynamics and there is no problem for higher-level agency to solve. For example, consider the convergence effect of increased gene flow between sub-populations, or consensus achieved via increased contact rate in other coordination games or consensus-finding scenarios (30). However, it is not the ease of finding consensus (under individual action) that we are interested in: It is the problem-solving competency of a cluster – the ability to enact specific collective action *despite* the presence of local optima that prevent such actions for the individual agents. Such collective action has the potential to collapse the effective dimensionality of the system dynamics, creating a new macro-scale process that searches combinations of module-configurations (in the same way that the micro-scale process searched combinations of individual units before the transition).

So, what would it take for a collective to self-organise such that it transitions from not having this basic competency to a new level of agency that does have this competency? How is membership in a collective identified bottom-up? What kind of relationships would cause them to act together, maximising long-term collective utility instead of short-term individual utility? And how do the necessary relationships arise given that relationships producing partial collective action are worse than none?

### Units, decision cycles and interactions

We explore the behaviour of individuals, units, that have a simple ability to decide an action (e.g., which side to drive on) and a ‘decision cycle’ that enables them to repeatedly return to an undecided state where they can decide again (Fig. 2). A Newtonian mechanics model, like that depicted by Waddington (31, 32)^3^ to represent developmental differentiation, is sufficient. Specifically, a ball rolling downhill and passing a saddle point causes it to branch left or right.

**Figure 2.**
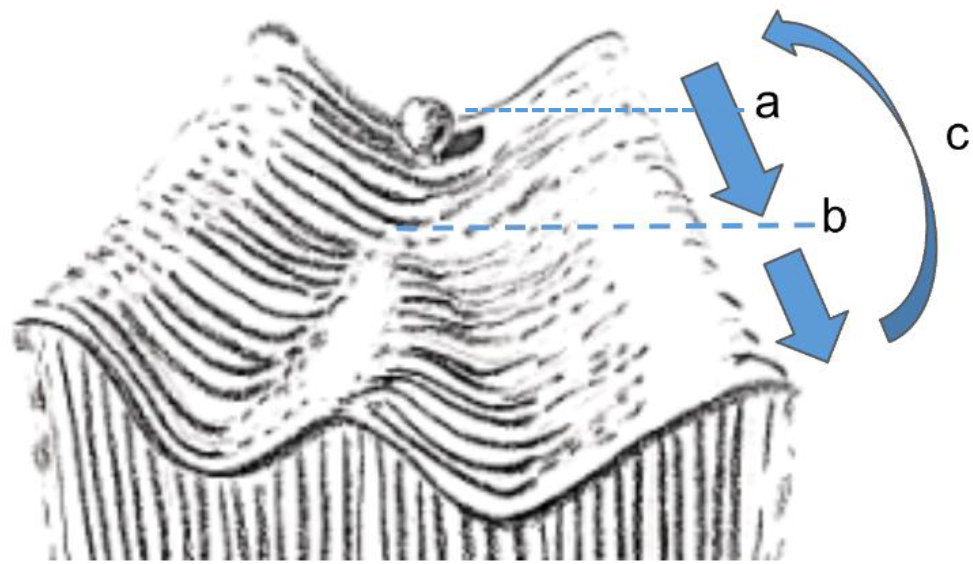
A single unit or primitive ‘decider’ modelled as a Newtonian system. In the ‘decision cycle’ of the unit, there is an amplification period which conceptually divides into two parts, an undecided or labile period (a), before the saddle point, when the unit is sensitive to small inputs but its outputs are weak; and a decided period (b), when the unit is no-longer sensitive to small inputs but its outputs (influence on others) is stronger. After these two parts of the amplification period, a new decision cycle is begun by reducing amplification and allowing the state to decay, i.e. effectively resetting the unit to the high-energy saddle point to begin again (c).

For each unit, *s*_*i*_, its state dynamics are described by:

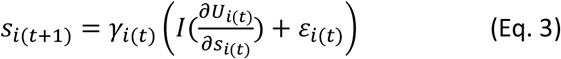

where *s*_(*t*)_ is the state of the unit at time *t*, γ_*i*(*t*)_ is a function controlling the period of amplification, *U*_*i*(*t*)_ is the utility of the unit, *I* is an amplification function, and ε_*i*(*t*)_ ∈ [-0.01,0.01] adds a small amount of noise to the amplification (drawn uniformly in the range). The decision cycle is modulated by γ_*i*(*t*)_ which oscillates sinusoidally. This controls whether the unit is in the amplification period (i.e. rolling down the hill) when γ_*i*(*t*)_ is high, or allowed to decay (back to the undecided state from where it can ‘decide again’), when γ_*i*(*t*)_ is low. Specifically, 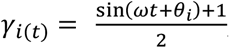, where *θ*_*i*_ is the phase of the unit’s decision cycle, and ω =0.01 is the frequency. Each unit thereby repeatedly takes one of two possible actions according to small differences in conditions at the decision point. The amplification function is defined as I(x) = max (min(100*x*, 1), −1). Each unit is essentially just an amplifier – a dynamical system that has sensitive dependence on initial conditions, with a decision cycle that repeatedly puts it back to this sensitive state (see Discussion for various biological interpretations of units and their decision cycles).

What is interesting about the model is not the individual units but rather the behaviour of many units in interaction with each other, and how the organisation of the timing relationships between them facilitates a transition in agency. This Newtonian model is adopted rather than an explicit selection-based or utility-maximisation model for two reasons: i) to make the point that nothing ‘clever’ is required on the part of an individual agent (i.e. its behaviours can be described by only a simple microscale equation of motion) and ii) because it is necessary that making a decision (rolling down the hill) is not an instantaneous event (an *argmax* model of utility maximisation is insufficient). Specifically, in the initial part of the decision-cycle the unit is not acting strongly (its output is not amplified) but it is sensitive to the influences of its inputs as it approaches the decision point. We can think of the inputs as tilting the decision surface slightly in favour of left or right. In the latter part of the decision cycle, after the bifurcation, the unit is no longer sensitive to its inputs (i.e. a slight tilt is not sufficient to move it from one basin to the other) but it is acting strongly (its outputs are amplified strongly enough to produce detectable ‘tilting’ influences on others) (see *a* and *b* in Fig. 2 and Fig. 3).

**Figure 3:**
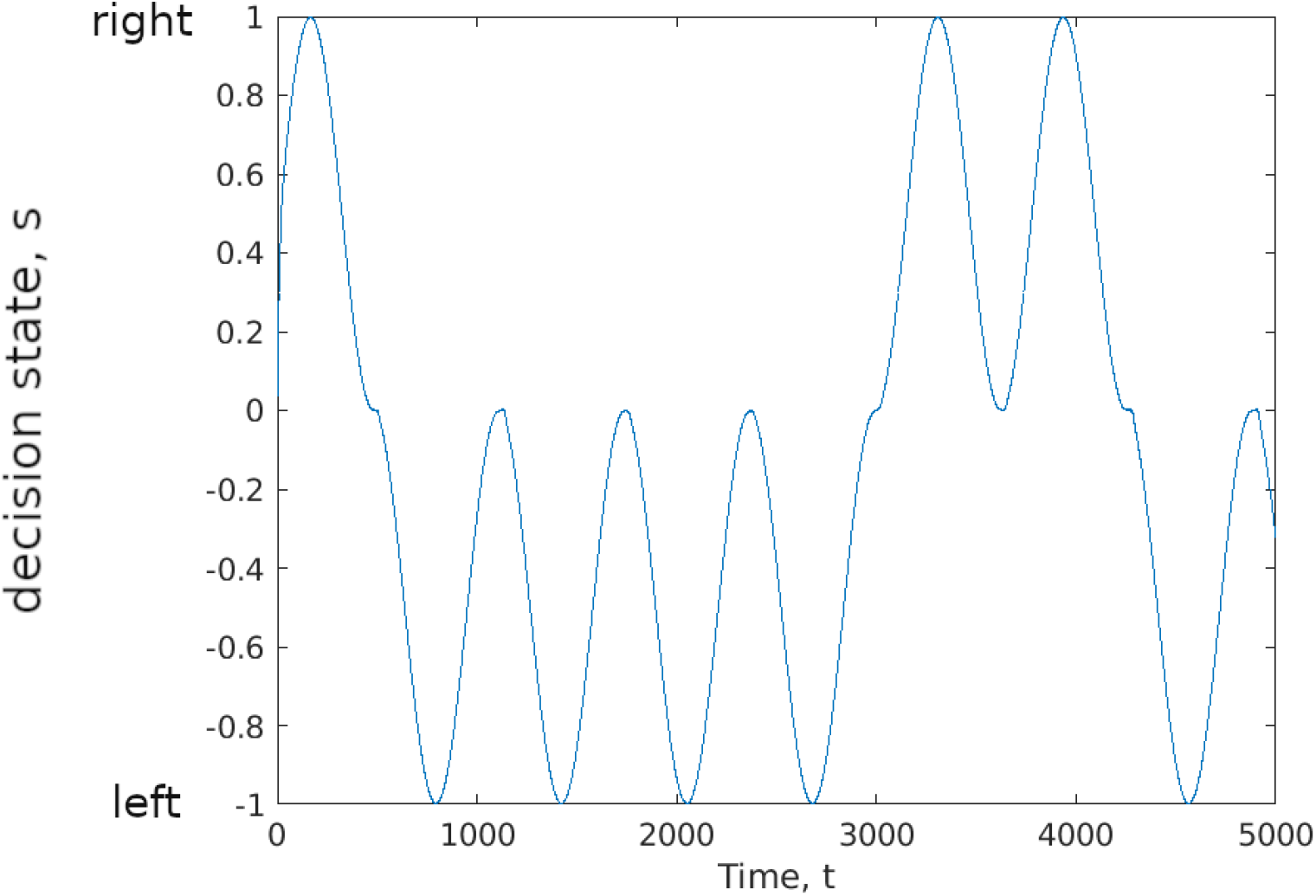
numerical simulation of a single unit (Eq. 3). A single unit has a ‘decision cycle’ which repeatedly pulls it back to the undecided saddle-point and then amplifies the state again, falling to either the left or right path (see Fig. 2). In this particular simulation there is only one unit, and thus due to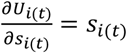, the left/right decision is arbitrary, determined only by the noise term. After the unit is decided, a small influence (or noise) is insufficient to change the decision. In the experiments that follow the decision is a response to the sum of inputs from other units (Eq. 4).

For multiple interacting units, where *c*_*ij*_ is the interaction strength of unit s_*j*_ on unit s_*i*_, the inputs to each unit involves a weighted sum of influences from all units (including itself):

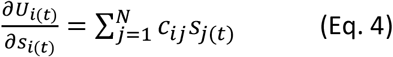

where variables are defined as per Eq. 3. Note that γ_*i*(*t*)_ continues to control the amplification/decay cycle as before, using the θ of that unit (initialised randomly for each unit, see below).

The back-and-fore between a state where an individual is sensitive to the world (but not acting) and a period where it is acting on the world (but not sensitive) is a key aspect of the model. This view of agency assumes that a system is either responding to its internal motives or to external influences and when it is doing one it cannot be doing the other. That is, being sensitive to one’s environment means necessarily that you are not entirely determined by your internal state; and taking action on the world means that you are driven by your internal state, to exactly the extent that you are not sensitive to external influences. Both sensitivity and action are necessary to be an agent, but simply adding up internal and external influences is insufficient for agential behaviour because this will lead to either to one overpowering the other or to an intermediate equilibrium of inaction where internal and external motives balance. Sometimes agency is modelled with the assumption that there are separate input (sensor) and output (effector) channels (22, 37, 38). But without presupposing such distinct one-way interfaces, and way for a system to be both sensitive and active is to move back and fore between taking (insensitive) action and being (inactively) sensitive. This allows for a system to be informed by world (reading the state of the environment and allowing its internal state to be changed by it) and to do work on the world (change the state of the world according to its internal state).

However, simply being able to repeatedly re-decide multiple times is not sufficient to solve the transitions problem. Specifically, if drivers make decisions in an asynchronous fashion, each driver will simply decide again to agree with the local consensus. The individual agency of the parts precludes the collective agency of the whole by holding it at a local equilibrium. This is what makes countries unable to take actions/to move to a different attractor.

### The hard problem of transitions

What kind of systemic change could produce a transition in the behaviour of the system so that countries become agential in the same way as the drivers they contain (i.e. able to choose between aggregate states that better-satisfy total stress)? One possibility is that if the components were arranged or organised differently then, given that their actions are context sensitive, they might make different decisions. This is the principle, for example, which enables an increased level of cooperation when interactions in social games are positively assorted, i.e. changing the interaction structure changes the incentives that an individual experiences (39). But, for a self-contained systematic model, we do not want to presuppose an interaction structure that produces collective action. Neither should an interaction structure depend on fortuitous happenstance provided by exogenous environmental factors (40). Rather it must be caused by the incentives of the individual components – they change their own interaction structure ‘bottom-up’ (41, 42).

However, if we suppose that individuals change their own interaction structure, then we have a new problem. If the changes to interaction structure that benefit long-term collective outcomes also benefit short-term individual outcomes, then it makes sense that individuals create those structures. However, in this case these structures align with what individuals are already doing, and the collective structure is explanatorily redundant (it doesn’t change behavioural outcomes) (9). Conversely, if the changes to interaction structure that benefit collective action are opposed by individual incentives (as is necessary for the collective to change behavioural outcomes), then individuals would not create that structure. And if neither of these is true, if changes to interaction structure are orthogonal to and hence have no effect on individual benefits, then again, there is no reason for individuals to create them (except by fortuitous happenstance). Is there any way out of this puzzle? How can individual incentives explain the creation of an interaction structure that ‘forces’ individuals to do something they don’t already want to do(20)? How can a transition be both systematic and meaningful (i.e. driven by bottom-up incentives and yet change behavioural outcomes)? We call this the hard problem of transitions in agency.

Here we explore one solution to this hard problem: namely, changes to interaction structures that control *when* individuals decide, not *what* they decide. In general, changes to timing do not directly control *which* actions individuals take (e.g. being caused to take action *A*, say, over action *B*), but simply modify *when* actions are taken (e.g. being caused to decide between *A* and *B* at time *t*_*1*_, say, over making this decision at time *t*_*2*_). Changes to the timing of individual decisions have the interesting characteristic that they can provide (small) immediate benefits to individuals without changing their decisions in that moment (i.e. they are aligned with current decisions) which means they are aligned with individual decisions and can arise systematically. Nonetheless, these changes do not commit individuals to their current decision in the long term either (because they do not control what the decision is at all), in fact, in a subtle way, they do the opposite and can thereby alter subsequent decisions in beneficial ways, as we will see. Thus changes to timing can be both systematic (because they are aligned with individual benefits at the time they are made) and meaningful with respect to altering behavioural outcomes (at other times).

### Entraining decision-cycles

If individual units are allowed to modify the timing of their decision cycles they will adjust them to locally maximise their utility or decrease stress. Since changes to timing are sensitive to combinations of behaviours and changes to behaviour are sensitive to differences in timing, this results in a simple form of positive feedback loosely analogous to the natural phase synchronisation, or entrainment, of weakly coupled oscillators (26).

Specifically, each unit, at a particular time *t*, has a phase, *θ*_*i*(*t*)_, that controls the timing of their decisions (Eq. 2). Initial values are drawn randomly, uniformly in the range *θ*_*i*(*t*=0)_ =[-π:π). The *θ* value of each unit is then incrementally adjusted on the following principle. Two units at time *t* that take actions s_i_ and s_j_ benefit from mutual synergy, e.g. due to taking mutually compatible actions, in proportion to their agreement with the interaction c_ij_ between them; specifically, c_ij_s_i_(t)s_j_(t). This quantity is positive if their signs are compatible with the sign of the interaction c_ij_, and its magnitude is increasing with the magnitude of each unit. Each state magnitude is a function of its phase, governed by γ_*i*(*t*)_ and *θ*_*i*(*t*)_. Over a given period *t*_*1*_ to *t*_*2*_, such as a complete decision cycle of a unit, the cumulative synergy is increasing with the integral of c_ij_s_i_(t)s_j_(t), and this is therefore affected by the phase difference between the states, |*θ*_*i*(*t*)_ − *θ*_*j*(*t*)_|. If two units have states that resolve the interaction between them (i.e. c_ij_s_i_(t)s_j_(t)>0), then the cumulative synergy available to a unit is increased when their phase difference is decreased, i.e. when they synchronise. A unit that can adjust its phase is therefore incentivised to adjust it toward other units with whom it is currently benefiting from mutual synergy in proportion to the magnitude of that synergy. The net movement in *θ* is determined by the total synergy with all other units at that time (integrated over a decision cycle). At a local optimum in the state dynamics, some pairs of units will have states that agree with the interaction between them, these attract values toward synchronisation in proportion to the magnitude of *c*_*ij*_. This principle is described by the following equation of motion (see Appendix A):

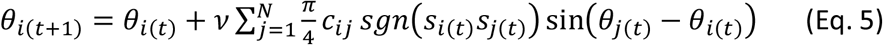

where *v* =3.10^−5^ is a constant controlling the rate of change of phase. Notice that this equation also allows for increases in phase difference (anti-synchronisation) when *c*_*ij*_ *s*_*i*(*t*)_*s*_*j*(*t*)_ < 0, so synchronisation only increases when states are compatible (i.e. their combination satisfies the interaction constraints they experience, on average over all the interactions they participate in). This incentive to synchronise is therefore not immediately apparent from the value of *c*_*ij*_ *per se* but requires units to first find an equilibrium under individual action (in the case where interactions are frustrated, establishing appropriate synchronisation may require sampling many such equilibria, as in previous work (43-47)).

Let us look more closely at the two-way interaction between the decisions that are made (*s* values) and when they are made (*θ* values) – or more exactly, between *combinations* of s values (correlated decisions) and *combinations* of *θ* values (phase differences). Recall that an agent must change back and fore between being sensitive (but inactive) and being active (but insensitive). This means that when agents adjust their timing to synchronise their activity, they also synchronise their inactivity – the moments when they are sensitive - and when agents are inactive and sensitive at the same time this has an interesting consequence. First, how does behaviour influence timing? If two units are already in agreement, synergy is maximised by synchronising their timing (this does not change their decisions at the time). The units that matter the most to changes in timing are the ones with the strongest agreement in s – in effect, *units that decide together wire together* – to imitate the Hebbian principle of associative learning seen in many networks with plastic interaction structure (48-53). In the other direction, how does timing influence behaviour? When a unit is undecided, it is not taking any action that cannot influence any other unit. If a pair of units act asynchronously then when one is sensitive it responds to the action of the other, and vice versa, which means their attention is on each other – or the memory of the collective state is held in these mutual and reciprocal influences. For example, I drive on the left because it was the best thing for me to do given what you were doing when I decided, and you drive on the left because it was the best thing for you to do given what I was doing when you decided. It is this circular or reciprocal state holding that prevents countries from being able to change their collective decision. Furthermore, this mutual amplification is most synergistic or beneficial to the participants when they are synchronised. But if synchronised, then a group of units are all sensitive (and inactive) at the same time, which means this mutual attention on each other is amplified at one part of the cycle and at the other side of the cycle this attention is momentarily broken – they have no influence on each other in this moment, and the memory is lost. In this moment of inner calm, the only remaining influence is from those that we are ***not*** synchronised with, i.e. the ‘out-group’. In short, synchronisation changes the ‘attention’ from themselves to their environment - from the ingroup to the outgroup. It is this that will allow the group to change its collective decision, as we will see. In short, the most important and reliably satisfied groups of states experience the strongest pull to synchronise, but whilst this amplifies the short-term benefit they receive from their mutual interactions, it also decreases their commitment to this decision; making it easier for them to escape the attractor created by their strong mutual influence.

### Evolving higher-level agency

The self-organisation dynamics described in Eq. 5 are relatively abstract but can be interpreted biologically. Here we aimed to test whether the same dynamics could be achieved by natural selection, in a simple setting. This evolutionary version of the model is implemented in the framework of adaptive dynamics. The system is composed of several populations (each corresponding to one unit) interacting with each other in the same coordination game described above. All the individuals within a population *i* have their own values for the heritable trait *θ*_*i*_, and the phenotype *s*_*i*(*t*)_ displayed by each individual is computed by Eq. 3 based on the value of this trait (the Left/Right decision is therefore a plastic phenotype, determined by environmental context, and the only thing that selection can do to influence this is to change the timing of the process). The fitness of each individual is given by Eq. 2 as the utility value, based on their phenotype and the average phenotypes of other populations. In other words, the influence on a given individual coming from other individuals (in the same population or some other) are modelled in an aggregated way for each population. Mutations on trait *θ* are considered under strong selection weak mutation assumptions such that they are rare enough that they happen one at a time, and the fixation or loss of each mutation occurs before the next mutation arises. The adaptive dynamics thus models each population as homogeneous, represented by an average *θ* value and an average state.

Given these simplifications, the evolution of the system can be modelled as follows. The initial state of the system is composed of N populations whose *θ* values are randomly drawn in [-π,π). Then, the simulation consists of iterations of mutation and selection on each population. During each iterate, the average phenotypes and fitnesses of populations are first computed with no change during 2π/ω timesteps, as a baseline to compare future changes in fitness due to mutation. Secondly, a focal population is randomly selected to be mutated. The mutated *θ*_*i*_ value is drawn from a normal distribution with mean *θ*_*i*_ and standard deviation σ. The average phenotypes and fitnesses are computed again during 2π/ω timesteps. If the new average fitness of the focal population is higher than its baseline, then it fixes in the population, and otherwise it is lost. With further mutation-selection iterations, populations undergo gradual change on their *θ* values according to local fitness increases.

## Experiments and results

### Basic dynamics of multiple coupled units (without entrainment)

Figure 4 illustrates the dynamics of interacting units (Eq. 4) with the modular constraints (Eq. 1).

**Figure 4.**
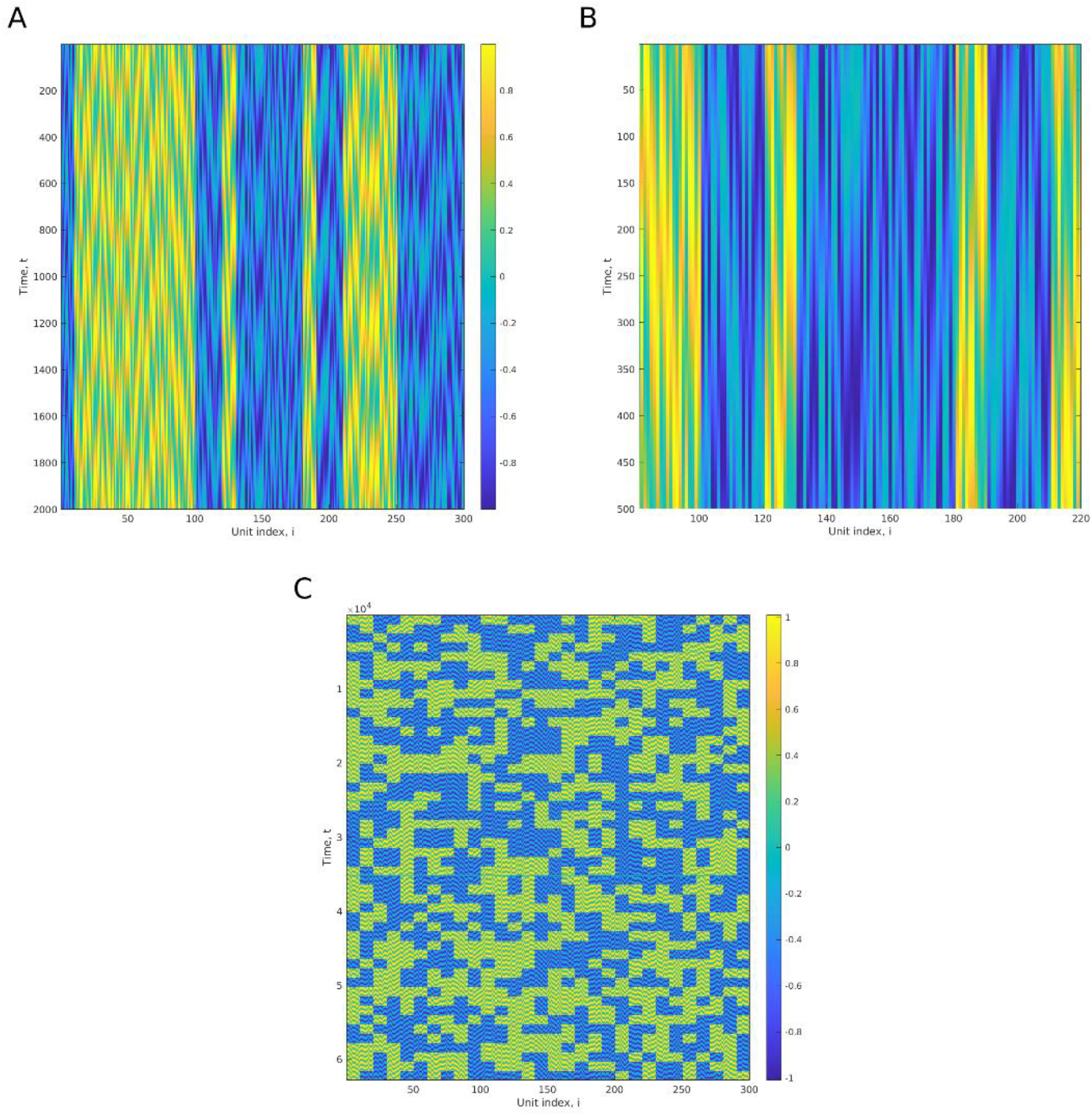
Simulation of 300 units with modular interaction constraints. (no entrainment). A) 300 units simulated for 2000 time steps from random initial conditions. Blue/yellow indicates the left/right decision of each unit, *s*_*i*_. Blocks of 10 units in the same module quickly adopt the same convention, either 10-left or 10-right. (Note that there is no spatial proximity of units; we have drawn units in the same module with neighbouring indexes (Eq. 1) so that we can see the stripes of 10 units easily in these figures, but their positions could be shuffled without altering the dynamics of the system). B) An enlarged section of 140 units for the first 500 time steps. At this resolution we can see more clearly the light blue ‘undecided’ periods where the states of the units are decayed close to zero (Fig. 3, Eq. 3), returning to their sensitive states. Notice that (without synchronisation) although they decide many times they always re-make the same decision since the influence of their neighbours has not changed. C) 50 independent simulations without entrainment, with different randomised initial conditions (at each multiple of t=1,250). No entrainment (random theta values). Since there are 2^30^ local optima under individual action, none of these runs is successful in finding a global optimum.

In Fig. 4 we see that the system falls to one of the 2^30^ possible attractor states, which are minima of Eq. 2. Each stripe of 10 units indicates that a subset of units has arrived at a configuration where they all agree with other units in the same module. But, in general, the likelihood that their decision agrees with some other module is approximately 0.5. Since there are 30 such modules, and 2^30^ minima, the probability of reaching the globally optimal attractor is very small (44, 47). In 50 independent simulations, the global optimum is not found (Fig. 4).

### Dynamics of coupled units with entrainment

In Fig. 5. we show a simulation of the system utilising the synchronising effect (Eq. 5). This shows how the system organises itself bottom-up (i.e. without assuming any processes of selection or reward above the level of individuals) in such a way that a specific collection of units becomes sensitive to and able to respond to its environment in a way that was not previously able (Fig. 5C, compared with Fig. 4A). Its behaviour is not just different, but more specifically a larger scale version of the original dynamics, i.e. before the transition, units responded and acted as individuals, after the transition, units respond as part of a single higher-level unit. Notice that the ‘waking-up’ of modules as units that can respond is quite sudden (and less than t=20,000 for most modules) (different collectives transition at different times, Fig.5.A, depending on the differences in their initial theta values, Fig. 5A). That is, the collective decides left/right in response to the pull of other collectives (Fig. 5D, white arrows), just like an individual unit (before the transition) decided left/right in response to the pull of other individual units.

**Figure 5.**
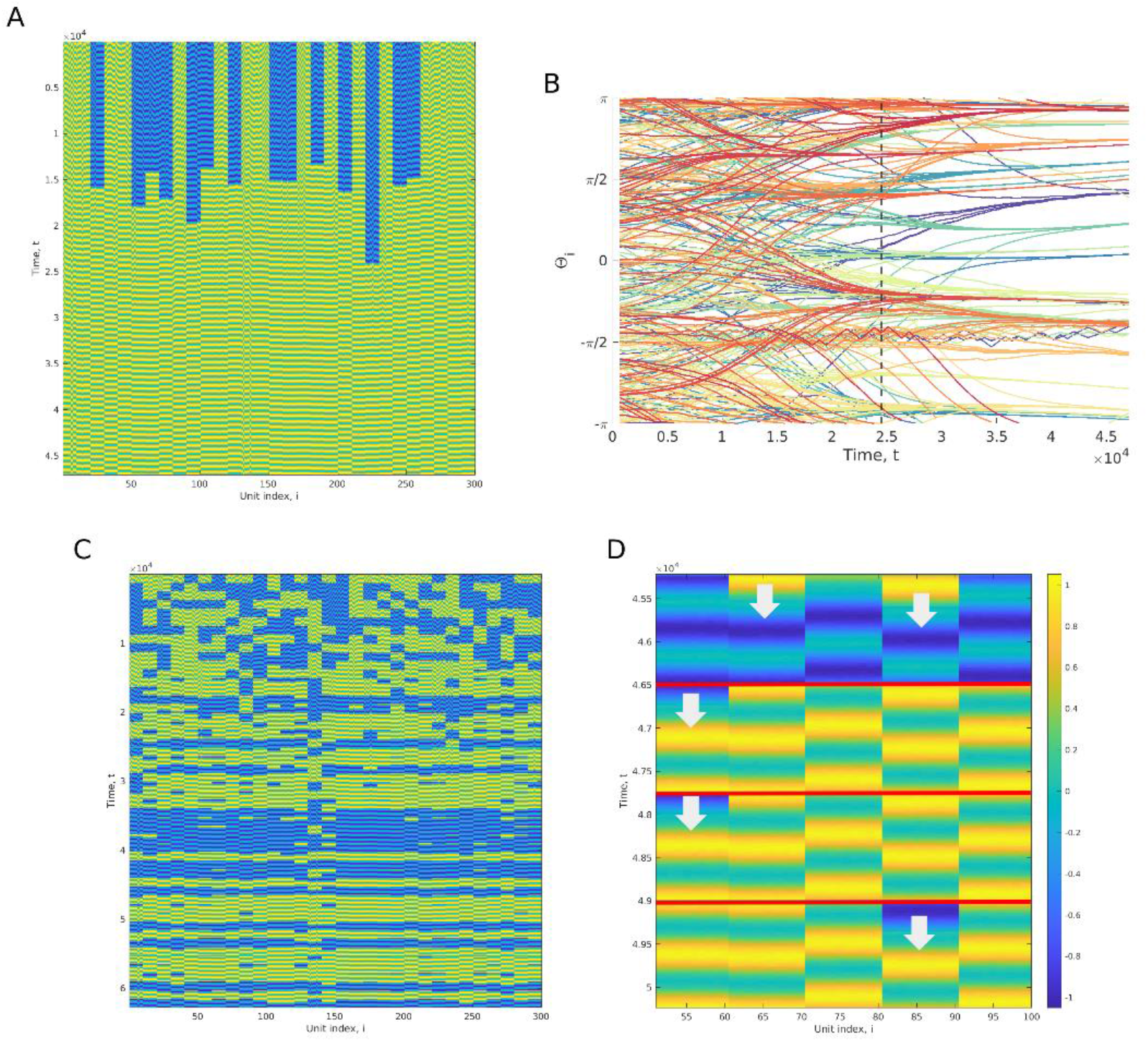
Simulation with entrainment (and multiple independent simulations after entrainment). A) With entrainment. Notice that at the start of the simulation the units are unable to coordinate between modules (as per Fig. 4). However, after around 20,000 timesteps, the system is fully coordinating all units across all modules. B) The evolution of theta values during the experiment in A. The thetas within a module are displayed in the same colour (some colours are reused for different modules). The vertical dotted line represents the time when modules have all become able to coordinate their decisions. C) Using the theta values found in A/B, we then perform 50 independent simulations with randomised initial states as before, but using the entrained theta values. These all find one of the two globally optimal solutions. D) A zoomed-in section from four runs in C, showing 50 of the units. Notice that the system still falls into stripes of 10 units very quickly, but such a block is able to change its decision (examples indicated with arrows), acting like a single macro-scale unit, responding to agree with the majority of other blocks.

This collective action causes the units to move through configurations that would be opposed by uncoordinated individual changes in state (e.g. a move from point a to point b in Fig. 1B). That is, when the members of the collective change one by one (or any less than half of them) they simply move back to the local optimum (as shown in Fig. 4B). Nonetheless, the collective action facilitated by synchronising their decision cycles causes them to move to a state where all of them are less stressed – not just a different equilibrium, but a better one where more of the constraints between units are satisfied. In other words, they exhibit collective action that facilitates long-term collective benefit that was previously opposed by short-term self-interest.

During the simulation shown in Fig. 5A (with entrainment) the theta values are being gradually adjusted (Fig. 5B). We observe that the theta values of a module change systematically in the direction of increased synchronisation even before a collective exhibits collective action (i.e. it is not motivated by the benefits of collective action it will later enable). To examine how changes in theta change the system dynamics, we extract the set of theta values at regular intervals of 1,300 timesteps during the simulation (Fig.5.A), and in a sequence of side-experiments, examine their effect on the ability of the system to find low energy states from random initial states. Fig. 6C. shows the energy of the system for each of these independent side-experiments. In comparison to Fig. 6B (which shows end-of-simulation energies from the independent runs without entrainment from Fig.4.C), this shows clearly how entrainment causes the system to get better at finding globally optimal states, not just locally optimal states.

**Figure 6:**
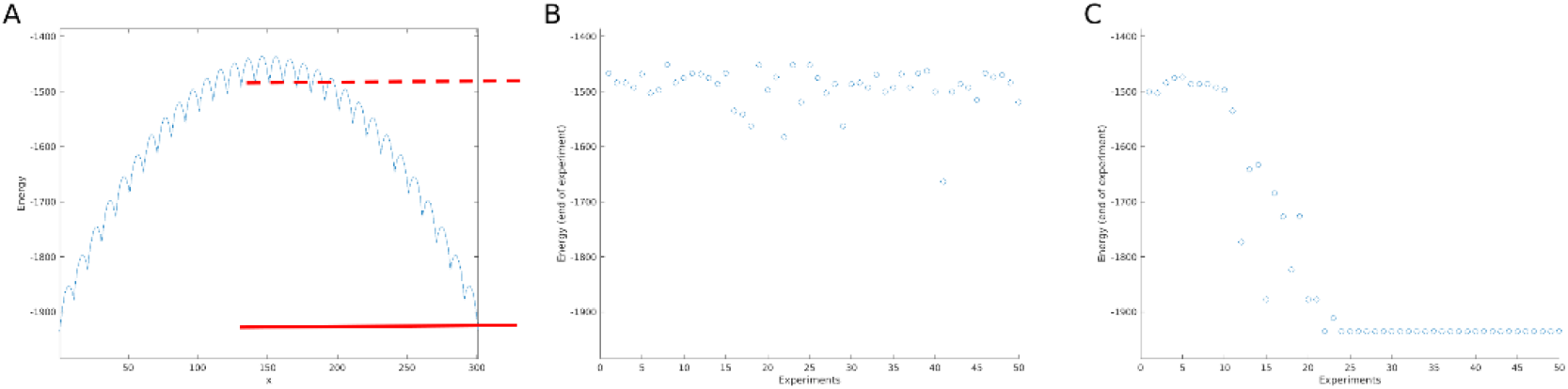
Energy surface of the system and energies of the attractors found without entrainment and as entrainment increases. A) A cross-section through the energy landscape (as per Fig. 1). The energy level of the most common local minima (where 50% of countries take one convention and 50% take the other) is indicated with a dotted red line. The energy level of the global minima is indicated with a solid red line. B) The energies that the system finds at the end of each independent simulation without entrainment (as per Fig. 4C). C) The energies that the system finds in side experiments (using samples of theta values taken from Fig.5.A). Notice that without self-organised synchronisation the system commonly finds the worst local optima (corresponding to the level of the dotted red line), and with self-organised synchronisation, after only about 20,000 timesteps, the system finds only the globally best local optima in every side experiment. This shows that the system has not merely found the global optimum, but found collectives that can reliably find the global optimum quickly from any initial state.

We then test whether the energy minimisation behaviour is merely a result of homogeneous synchronisation, or a result of organised and specific synchronisation (i.e. within modules only). Fig. 7A shows the behaviour of the system in 50 independent simulations when the synchronisation is system-wide, global across all the units in all the countries. This does not exhibit transitions or the increased energy minimisation behaviour.

**Figure 7:**
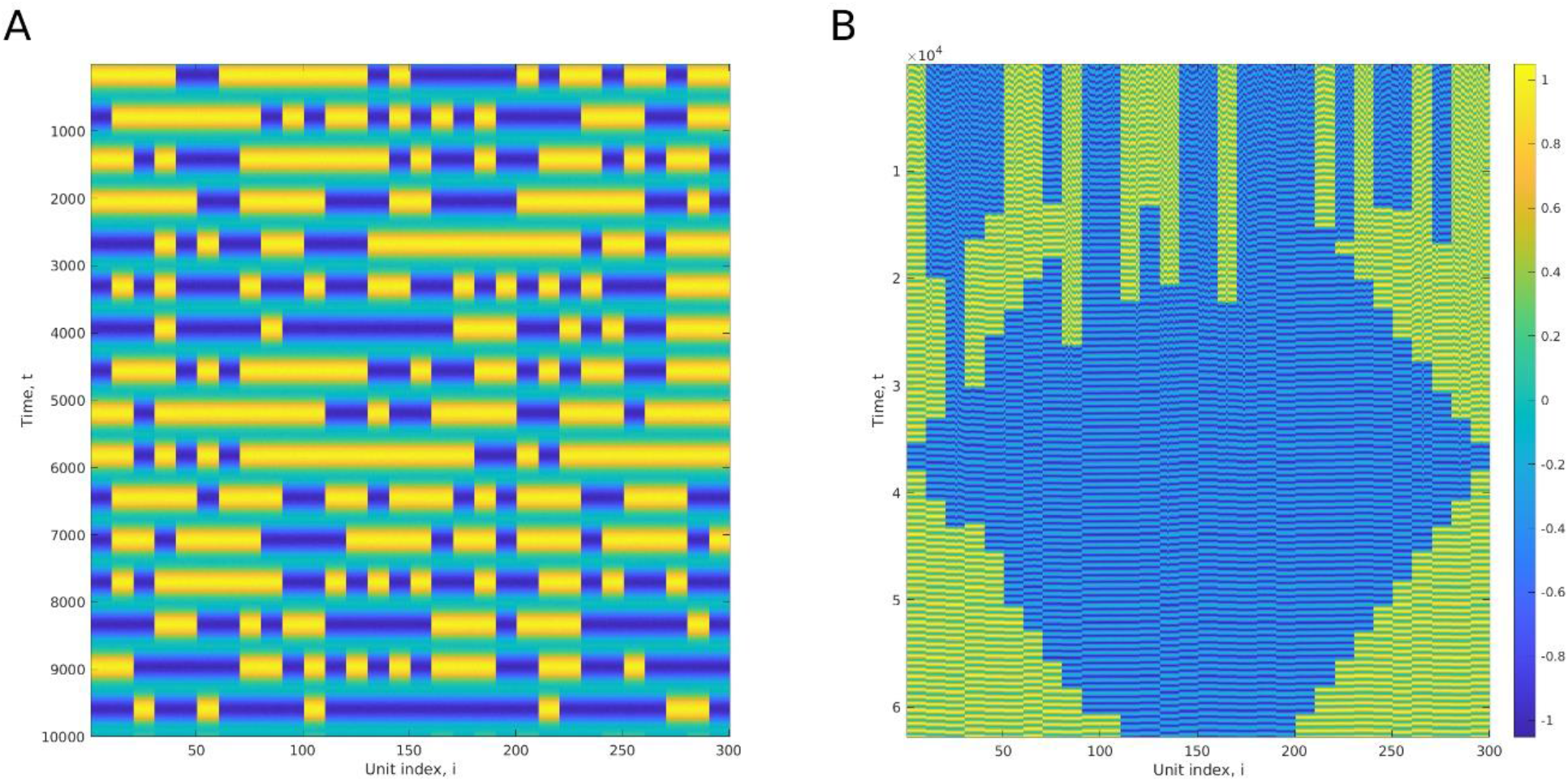
Global synchronisation does not facilitate global energy minimisation, specific synchronisation enables global energy minimisation in a sequence of different problems. A) Behaviour of the system when the synchronisation is global, i.e. across all the units in all the countries (no self-organising entrainment). Note that this does not resolve conflicts between countries. Specifically-organised synchronisation is required to enable one country to respond to another. B) In contrast, the specifically structured synchronisation emerging from entrainment dynamics, results in agency that can not only find configurations of globally minimal energy in the original problem, but can do so even in a ‘moving target’ problem. Here the dynamics of the units after entrainment is exposed to a sequence of different environments (alternative constraint problems). Specifically, rather than a simple game of global coordination, the game used here rewards a middle section of countries for disagreeing with the remainder, and the size of the middle section increases and decreases over time. This shows that the modules can be identified from a non-stationary problem, so long as the within-module interactions are consistent, and more importantly, they not only enable effective stress-reducing responses for the problem in which they originated, but for any problem based on these modules (i.e. that can be solved by following an energy gradient at the macro-scale).

With entrainment, the ability of the system to find the global optimum does not derive from having learned the specific location of the global optimum, nor from merely global synchronisation of all units. It has not learned the solution; rather, it has learned how to move in a way that finds solutions. And it has not learned how to find solutions to one particular problem but to the class of problems with this module structure, i.e. solvable through some combination of these collective movements. The specific collective action that results can solve any problem that involves solutions made from combinations of these modular constraints (by hill-climbing in modules, as if enacting a macro-scale dynamics). Accordingly, if the environment rewards a different combination of modules, or the problem changes over time, agents can respond appropriately, tracking a non-stationary problem (Fig. 7B).

### Evolutionary dynamics of coupled units

Entrainment occurs in entities that can adapt to external constraints either as strategic utility-maximising agents or through evolutionary adaptation. Here we examine the synchronisation dynamics described in Eq. 5, and their effect on the coordination of states, under natural selection. Considering that each unit is a population of organisms in which theta values are genetically encoded traits, in Fig. 8 we show a simulation for a system of 7 modules of 7 populations (i.e. 49 populations in total) with individuals evolving these traits to maximize their fitness (minimize their energy) with an adaptive dynamics model. This shows that the system does not require deterministic dynamics (such as provided by Eq. 5) in order to organise itself bottom-up. Starting from random mutations only, variation and selection is sufficient for the system to reach the same level of synchronisation of theta values in each module (comparing Fig. 8A with Fig. 5B). In turn, as theta values converge within each module, modules become able to respond to external constraints (see Fig. 8B). As per Eq. 5, this phenomenon only requires the short-sighted maximisation of fitness.

**Figure 8:**
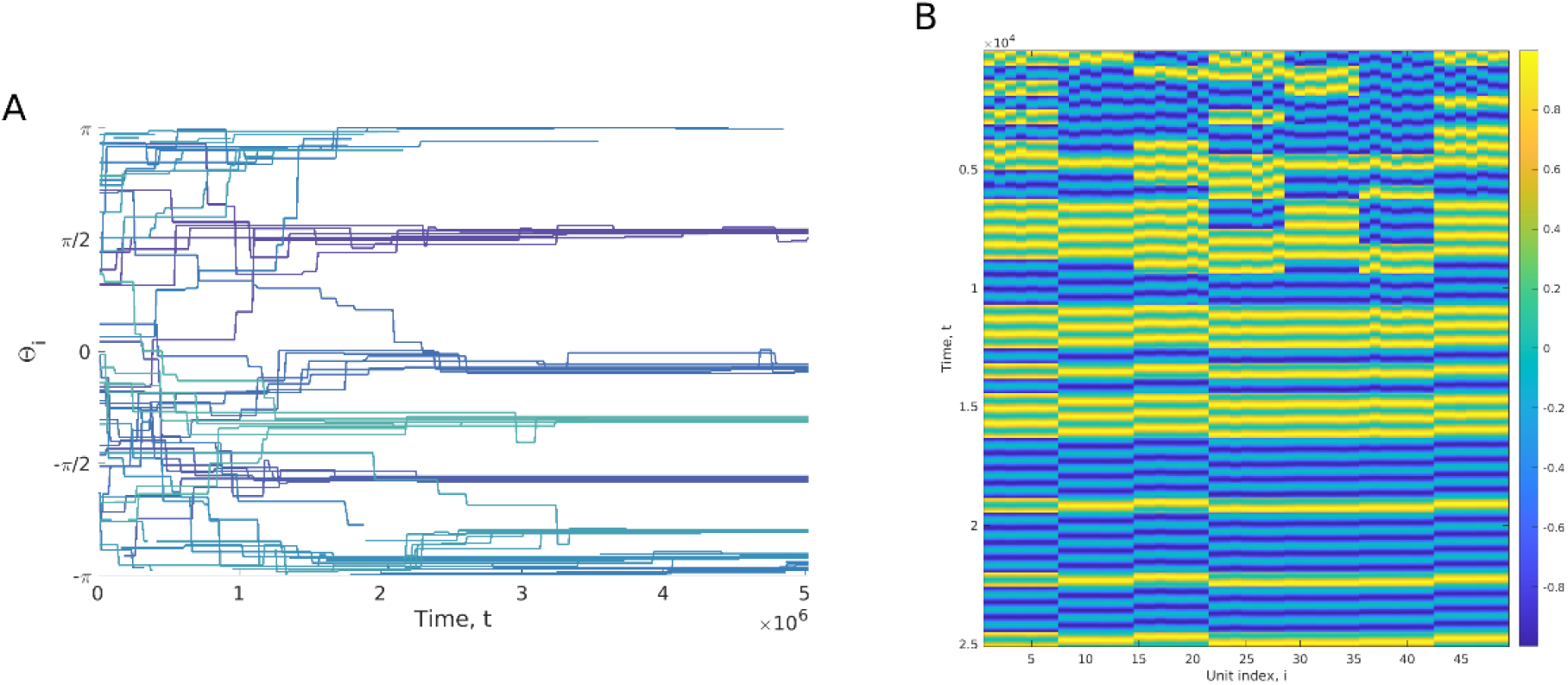
Specific synchronisation can be achieved by a process of natural selection and yield the ability for modules to reach the global fitness optimum. A) The evolution of average theta values during the adaptive dynamics experiment for a system composed of 7 modules of 7 populations. With sufficiently large mutations (σ=0.5), the populations are able to adapt to reach synchronisation within each module by natural selection. The thetas within a module are displayed in the same colour. B) The evolution of average expressed phenotypes (states) due to the change in theta values displayed in A). The colors displayed here do not represent continuous change, but rather 40 mutation-selection cycles recorded every 50 cycles over the course of the simulation. As in the other simulations, populations are initlally able to coordinate their phenotypes within each module, but not across modules. As the theta values evolve by natural selection towards within-module synchronisation, populations become able to fully coordinate the phenotypes of all populations within all modules.

Mutations that decrease the difference of theta values between populations evolve because it is of immediate benefit to express the same state at the same time as other strongly-interacting populations, not because they will, in the future, enable a set of populations to change state together. This short-term gain and the consequent long-term gain provided by collective action are caused by the same changes in phase. But this is not a coincidence because they have a common cause; specifically, strong interactions confer large benefits for units that synchronise and it is the same strong interactions that created the multiple local equilibria in the state dynamics. Groups of these strong interactions hold the attention of the units they contain until such a time as they synchronise and their attention thus turns outwards to other groups (54).

Notice that the change in theta values is not as smooth as the dynamics of Eq. 5 (Fig 5B), due to the stochastic nature of mutations. This has the consequence of increasing the time required to reach synchronisation by several orders of magnitude (10^4^ timesteps for deterministic change of Eq. 5 and 10^6^ for stochastic evolutionary change). This time increase arises even though the system studied here is simpler (only 7×7 populations, compared to 30×10 units) and the evolutionary algorithm applied here (adaptive dynamics) ignores the effect of genetic drift, migration, and recombination. This result shows that the emergence of higher-order behaviour by natural selection is possible by the same principles, albeit less efficiently.

### Dynamics of hierarchically coupled units with entrainment

We showed above that gradual change in the timing of decisions of units within a module could lead them to transition from acting as individuals to acting as parts of a single higher-level unit. Here we explore whether this continues to scale: i.e. can gradual change in the timing of decisions of higher-level units (i.e. modules) lead them to transition towards acting as parts of a single even-higher-level meta-module? In order to test this, we utilise a hierarchical version of the modular driving conventions problem. The interaction strengths are now defined as:

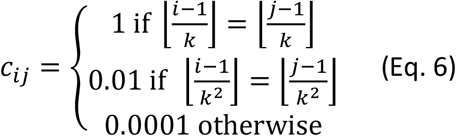

where i and jϵ{1,…N} are the indexes of the pairwise constraints over N variables, ⌊x⌋ is the floor of the value x, k is the number of units in each module, and k^2^ is the number of units in each meta-module.

Fig. 9 shows the dynamics of states over time with entrainment using Eq. 5, using N=343 and k=7. As observed in Fig. 5 for the non-hierarchical modular problem, units are first unable to coordinate their states across different modules, then by synchronising their decision cycles, they acquire the sensitivity required for modules to respond to the pull other modules. Because the problem is now hierarchical, further synchronisation between units of a meta-module can also confer them a new level of sensitivity, making the meta-modules able to respond to the pull of other meta-modules. In that sense, it shows that the transition observed at one level operates through multiple levels; transitioning through successively higher levels of deciding and collective action.

**Figure 9:**
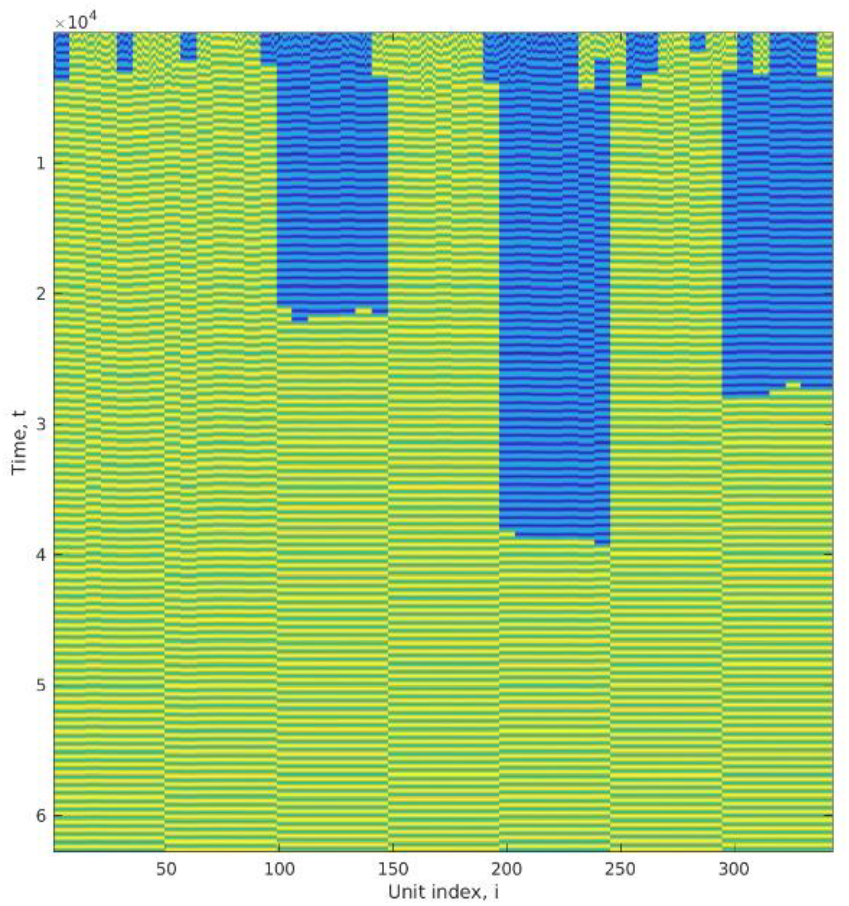
Synchronisation can be achieved hierarchically. This simulation displays the dynamics of 7 meta-modules of 7 modules of 7 units to solve the hierachical modular driving conventions problem defined in Eq. 6, with entrainment following Eq. 5. Units are first able to coordinate their states within each module, before modules become able to coordinate their states, until meta-modules finally become able to coordinate their states after around 40,000 timesteps. Notice that here ν=3.10^−4^, that is 10 times higher than in the modular driving conventions problem.

However, with σ=3.10^−5^ in the simulation in Fig. 5, the ‘waking-up’ of modules took only 20,000 timesteps to occur. In Fig. 9 with a rate of change in theta values 10 times higher (σ=3.10^−4^), the ‘waking-up’ of meta-modules does not occur until 40,000 timesteps. If the transition was truly scale invariant, one could expect the transition from units to modules at the same rate as modules to meta-modules. This is not the case because the synchronisation of theta values between modules of a same meta-module requires the theta values of all the units involved to synchronise. Units can become able to respond to their environment as parts of several orders of higher-level units, but in this model, higher-level modules do not have the ability to respond to selection as a single unit.

## Discussion

The experiments illustrate a simple abstract model of a transition in agency using behaviour synchronisation. This shows how a very simple agential competency – deciding among actions to maximise individual utility or minimise stress – arises at the collective level. The consequences of this transition is that it originates a problem-solving competency at a new scale of organisation, and we demonstrate that this involves a collective action that moves against (unilateral) individual utility in order to arrive at improved collective utility in the long term. How is this possible?

### Timing and the apparent paradox of agency

Higher-level agency carries an apparent contradiction between a microscale behaviour that ‘fully describes’ the system and a macroscale behaviour that somehow ‘over-rules’ the microscale behaviour. For example, if I merely do what my cells are already inclined to do then I, as a putative higher agent, am explanatorily redundant. In order to be an agent that is not redundant, I have to make my cells do something that over-rules their individual inclinations – change behavioural outcomes. Nonetheless, we suppose, I am nothing more than the collective action of my cells. So how can this be possible?-Our model helps us think about this apparent contradiction and suggests that changes to timing might be a key mechanism to mediate between micro- and macro-scale processes.

On the one hand, the inner workings of the model are described at the microscale in a reductionist way. The higher-level agent is not a variable in any of our equations of motion, and in this sense it is causally redundant. On the other hand, the behaviour of the microscale components after the transition affords this apparently goal-directed behaviour at the collective level, i.e. it moves against individual utility in the short term in order to arrive at improved collective utility in the long term. This new organisation (the changes in timing) is not causally redundant, i.e. it alters outcomes, by creating a new locus of sensitivity and control that was not previously present at that scale (7, 19, 55, 56).

Of course, changes in timing are also part of the microscale dynamics, and with one kind of timing, the same individuals with the same competencies may exhibit quite different behavioural outcomes than they do with another kind of timing (57, 58). Specifically, in our model, when all the units within a collective are in their sensitive state at the same time, none of them is exerting strong forces on the others. In this moment, they each can be influenced by the weak forces external to the collective (from other units or collectives). At the macroscale, the collective effectively moves from one local optimum to another following the macroscale energy gradient (Fig. 1.B, broken red curve), even though all intermediate configurations have higher energy (Fig. 1.B, solid red curve). The direction of change in these collective movements is not stochastic but goal-directed, i.e. toward configurations with lower collective stress, even though it involves a set of changes that, if performed unilaterally, would be opposed by individual stress.

What is interesting about timing therefore is that it does not merely alter when you are sensitive and when you are not, but through synchronisation, it alters your *attention*, i.e. who you are sensitive to and who you are not sensitive to. But how do individuals end-up attending more to weak external interactions and less to their strong internal interactions? This is because the decision process cycles between being sensitive (but not acting) and acting (but not sensitive). Accordingly, a mutual incentive for strongly dependent units to act at the same time, also results in a moment when these strongly dependent units do not pay any attention to each other. This effects a change in attention from ‘inside’ the collective to ‘outside’ – not by paying more attention to the weak external signals but by momentarily paying less attention to internal ‘business as usual’. This synchronisation of behaviours within the collective, i.e. *inner alignment*, thus creates an ability to respond to external influences at a new level of organisation.

Moreover, synchronisation does not change the behaviour of the parts (e.g. choosing between left and right), but it changes how the behaviour of the parts is organized, which allows the parts to do better at what they normally do given the constraints between them. Regulating the attention of the parts does not take away their agency from them, but it gives them knowledge about the best decisions they can make. This knowledge is *internal* to the system in the sense that it belongs properly to each module (the phase differences between its components) and it is not superimposed by external conditions. A level of organization with internal states which allow the parts to forgo short-term benefits in order to reach greater long-term gains can be qualified as a new level of agency (20). This phenomenon can scale up so units that have already transitioned to a new level of agency can transition again to an even-higher level of organization (see Fig. 9 and the hierarchical modular problem). Here units in our model are able to leverage higher-level agency (i.e. higher-order local equilibria) to reach the global equilibrium. It is debatable whether this ability constitutes an even-higher level of agency, but it is clear that scaling-up transitions does not necessarily mean that the higher-level units themselves are able to transition. Transitions are possible because there are traits that can be organized to produce internal states. Thus, any further transition would require higher-order traits (e.g. meta-phase differences, as a phase for higher-order oscillations) that can be organized to produce the even-higher level of agency.

True energy-minimisation processes do not go uphill, of course. The energy function over discrete states described by Eq.2 (and Fig. 1) ignores the ‘in between’ states of the continuous decision cycle – in particular, the high-energy state occurring when a unit is reset to the saddle point. When the behaviours of units are uncoordinated this is a reasonable simplification because the high-energy states are effectively lost in noise (Fig. 5). However, as their behaviour becomes synchronised, these high-energy states enable the units to escape from the apparent local optimum they are otherwise trapped in. Thus, properly understood, the collective action behaviour of the system does not take it through states that are higher in energy. It is essentially an entropic pressure that keeps disorganised collectives at local minima, and escaping this is an organisational or informational issue, rather than a true energy barrier.

### Associative learning and decentralised positive feedback on correlated behaviour

This information gain, necessary for a transition, is possible because, through synchronisation, the units are ‘learning’ something about the structure of the dependencies in the problem (44). It is this information, acquired bottom-up through natural distributed processes (without a higher-level selection process or utility-maximisation principle at the collective level), which enables a collective to overcome the apparent energy barrier created by disorganised internal behaviour.

The entrainment of weakly-coupled oscillators, causing synchronisation or phase locking (26, 59), is a ubiquitous physical phenomenon that spontaneously organises the timing of repetitive behaviours (e.g. from pendulum clocks, to fireflies, to pedestrians) (60, 61). Unlike previous models, however, it is not the synchronisation *per se* that constitutes the macroscale behaviour we are interested in (e.g. synchronous flashing of fireflies). Rather we are interested in how synchronisation enables different behavioural outcomes, i.e. in this case, resolution of the constraints between the Left/Right state variables.

The synchronisation of units is related to a principle of positive feedback on correlations; the more two things coordinate, the more the relationship between them changes in the direction that makes them more coordinated (49, 62). This positive feedback on correlations is an analogue of a simple unsupervised associative learning principle, or Hebbian learning principle (i.e. ‘neurons that fire together wire together’) (63). This does not require any sophisticated learning machinery or an external teacher, reward signal or system-level selection process. It occurs spontaneously in dynamical systems described by networks of viscoelastic connections, i.e. connections that give-way or deform slightly under stress (62). In many kinds of networks this kind of positive feedback is a natural consequence of *individual-level* natural selection acting on heritable variation in the relationships with other individuals – and as modelled here, a natural behaviour of coupled oscillators. The fundamental principle is that state variables change to minimise stress given the current structural variables, and structural variables change to minimise stress given the current state variables. Previous work has demonstrated conditions where this organises relationships between components in a manner that produces non-trivial collective adaptation without presupposing a collective-level selection process (49-52, 64). However, that work did not demonstrate a transition in the level of agency in the sense shown here. Here, turning the positive feedback principle to act on timing, and thus attention, produces a subtle but crucial shift in scale, i.e.: the more often a subset of individuals (with their individual attention on each other) exhibits synergistically coordinated actions (and the more they synchronise) the more such a *collective* (with its collective attention outwards) is able to find synergistically coordinated actions *with other collectives*.

### Transitions in evolution and development

An individual unit could represent many different types of biological ‘decision processes’ including cell differentiation (given a set of morphogens produced by other cells), a genetic selection process (given a set of epistatic fitness interactions with other genes), an ecological selection scenario (given community interactions among species), or gene-expression dynamics (given regulatory interactions between gene expression levels). In many such domains, a modularly weighted coordination game can be used as a fulcrum to think about multi-scale agency and multi-scale optimisation (44, 46, 47, 64, 65). For example, in developmental systems, a similar scenario may obtain for a tissue where each cell has a phenotype, such as planar polarity, that needs to be coordinated with neighbouring cells through local communication. In a genetic evolutionary context, there is a deep homology between the transitions in agency problem (maximising long-term collective fitness contra short-term self-interest), and the evolutionary problem of crossing valleys in a fitness landscape (with a natural selection process that can only respond to the average excess of individual alleles) (66-68).

In the evolutionary transitions in individuality, a central problem is the conceptual switch from comparing the fitness of individuals to comparing the fitness of collectives (a.k.a. “fitness-1 vs fitness-2” (9)). The crucial significance of our model for this problem is that it is a bottom-up model: The transition occurs without any selection among alternative macrostates, such as a higher-level selection or utility-maximisation process that compares or competes one collective state with another. Nonetheless, the model suggests that if individual selection can act on reproductive timing (Fig. 8) then a system can behave *as if* it follows an explicit higher-level selection process – insomuch as it facilitates movement from one collective state to another superior collective state, apparently over-ruling individual-level interests (i.e. as if fitness-2 is over-ruling fitness-1).

Evolutionary transitions often involve control over the reproduction of components (suppressing fitness differences within the collective) and control over the production of phenotypic variation at the group level (creating fitness differences between collectives) (9, 10, 13, 29). A possible analogy is that a decision-cycle corresponds to the reproductive life-cycle of lower-level units, and synchronisation corresponds to cell-cycle synchronisation (e.g. of initially undifferentiated cells in early embryogenesis), followed by development of a collective phenotype (exhibiting a new collective sensitivity to initial conditions). Intuitively, having a developmental process that repeatedly transforms a mass of (simultaneously undifferentiated) embryonic cells into a differentiated adult phenotype is an intrinsic part of what distinguishes an organism from an ecology of individual cells (10). Developmental processes within multi-cellular organisms, for example, are a result of transitions in individuality. These constitute two-levels of variation and selection (at the somatic level of cell-lineages within a lifetime and at the level of the organism over multiple generations) one nested inside the other, both in terms of organisational scales and temporal scales. The ability of development to explore phenotypic options within lifetime is a particular example of two-level variation and selection process characteristic of exploratory mechanisms, and playing a central role in evolvability (69, 70). Note that when the cells of the initial developing embryo are in an undifferentiated pluripotent state at the same time, the organism is exquisitely poised at a saddle point in its developmental dynamics – able to differentiate in different ways responsive to small signals. This changes the kind of effect that genetic mutations can have, moving the phenotypic response to mutations, and hence the selective response, to a higher level of organisation – e.g. controlling traits that affect the complementarity of cell types (e.g. cell communication and phenotypic plasticity) rather than controlling their static fitness-affecting traits (10).

The emphasis on timing potentially alleviates the problem of why individuals would evolve relationships that cause them to cede control over their own reproduction (9). Whilst it might be tempting to appeal to the long-term collective benefit thereby achieved, remember that a collective is not a thing that has a fitness until *after* the transition, and relationships that cause only partial changes in behaviour are worse than none. A focus on timing alleviates this because, before the transition, changing the synchronisation of behaviours with strongly interacting units is individually beneficial given the actions they are already doing (fitness-1). Later, when all the units of a collective are in synch with each other, they can effect a collective change to a new behaviour, providing a fitness benefit to all members (effectively fitness-2). The short-term (proximal) benefit of synchronising and the long-term (ultimate) fitness consequences are therefore aligned but not the same. The proximal causes of such synchronisation may be various and is common across many biological systems including synchronisation in development (13). Reproductive synchronisation in particular (e.g. synchronised flowering, spawning, mating, etc.) is common in many organisms and fundamental to the equalisation of within-collective fitness characteristic of evolutionary transitions (71). Nonetheless, in this model it is not a coincidence that the structural relationships relevant to short-term benefit are the same interactions relevant to long-term benefits; They have an underlying common cause (54). Specifically, short-term benefits come from synchronisation of actions at the current local equilibrium and long-term benefits, requiring escape from this equilibrium, come from ignoring these interactions and turning attention toward others. These are different benefits on different timescales, and only the former can be the reason that a particular pattern of synchronisation occurs. Yet, it is non-coincidentally the same interactions involved in both benefits. The twist in the story is that – given the constant movement of the decision cycle between (insensitive) action and (inactive) sensitivity - attending to the interactions that are most important, and synchronising decisions with them, also creates a moment when these interactions are the least important and attention turns elsewhere.

### Doing without choice

Agential behaviour is often associated with a notion of choice (72) – as though a system *could have done otherwise*. This introduces notions of ‘deliberating’ among future possibilities, such as explicitly enumerating alternatives and selecting among them (73). A classical *argmax* model of a utility-maximising agent, for example, is an explicit enumeration of possible actions and their consequences. Likewise, in a natural selection process, an entity is duplicated with variation, developed and the phenotypic outcomes naturally subject to differential survival and reproduction. In this paper we specifically excluded a mechanism of explicit deliberation; a mechanistic model of transition in agency must utilise only a microscale equation of motion and do without an explicit process that compares the fitness or utility of one macrostate to another. Whilst a process that enumerates possibilities and selects among them can be physicalized in principle (e.g. evolution by natural selection is not unphysical), a model of ETIs cannot assume this. It is important, therefore that nowhere in our model is there any enumeration of future possible states or comparison between them. Nonetheless, something that appears to be ‘deliberate’ (i.e. a movement toward a specific long-term outcome) can arise without presupposing an explicitly deliberative process. The appropriately synchronised collective is simply able to ‘roll downhill’ following the macroscale energy gradient. This has outcomes equivalent to deliberation or equilibrium selection, but without involving a higher-level selection or higher-level utility-maximisation process.

Notice that a reductionistic perspective does not recognise any need to ascribe top-down causation. Individuals comply to individual incentives; They are not mysteriously forced to do something they do not want to do, either before or after the transition. Instead, they do different behaviours simply because the situation they are in after the transition is different from the situation they are in before the transition. Moreover, the causes that explain the change in the situation (the organisation of the phase differences) are also determined by individual-level actions. Whilst a top-down perspective recognises that the organisation is causing different outcomes, and the organisation is not a property of any individual part (by definition), the reductionist only sees this as a temporal synergy of individual behaviours that could not be otherwise given the microscale entities and their interactions as they arise in this moment. This is just to say that a reductionist perspective is one that does not recognise organisations as causes (17, 18, 20, 56). This is a self-consistent position; however, note that organisations can be under-determined by microscale causes, in particular when there is strongly divergent dynamics followed by strongly positive feedback; for example, when a learning system converges on a particular way of solving a non-linearly separable function. This means that the organisation depends on “the history of the system, the whole system, and nothing but the system” and without this history the predictive power of microscale causes is minimal (20).

### Toward more sophisticated agents

Whilst this model forms collectives with persistent behavioural competencies, there is no sense of an agent that maintains physical boundaries in space, controls an internal metabolism or exhibits any complex behavioural repertoire. The new level of agency also has no internal computation or information integration. The latter, although limiting, is also a point of interest. That is, units act together, not because they receive useful signals from one another, but because they stop receiving strong self-reinforcing inputs from one another that overpower their sensitivity to external inputs. Nonetheless, in some other scenario, if their experiences of the external context were localised or otherwise non-uniform, coherent collective action would require internal information integration (10). Perhaps, though, these sorts of refinements first require a change in the level of agency provided by the simple sort of collective benefit demonstrated here.

## Conclusions

After the transition, the collectives that emerge in this model show a new level of sensitivity that they did not have before the transition – an ability to decide between two collective states to maximise collective utility. Thus they exhibit agency in the same (limited) sense as the individual parts did. The formation of the collective is systematic because it is caused (‘bottom-up’) by immediate short-term benefits to the individuals involved – without imposing a suitable organisation or assuming a fortuitous change in exogenous conditions to initialise one. The collective is nonetheless meaningful because it changes outcomes – it shows ‘top-down’ causation insomuch as it creates a situation in which individuals therein change their decisions in a way that would be opposed by unilateral (unsynchronised) action. The resulting behaviour appears goal-directed insomuch as the collective moves systematically in the direction of collective benefit in advance of receiving it (i.e. without sampling or selection of alternative possibilities). Accordingly, we suggest that this model provides a worked example of a system that solves the hard problem of transitions in agency – a transition that is both meaningful and systematic – and that this kind of synchronising behaviour, common to many physical systems, may be key to understanding transitions in other systems.

## Acknowledgements

Thank you to David Prosser for comments on the manuscript, and Jonathan Young and Lucas Mathieu for conceptual discussion. The authors gratefully acknowledge the support of grants 62230 (RW, TT), 62220 (RW) and 62212 (ML), from the John Templeton Foundation, and BBRSC grant BB/P022197/1 (CB).

## Appendix A: Optimisation of interaction synergy by changing phase difference between units

Following the section *Entertaining decision cycles*, we consider that units will modify their *θ*_*i*_ values to increase their synergy with other units. Indeed, if the utility of their interaction (u_ij_(t)=c_ij_s_i_(t)s_j_(t)) is positive, then they should decrease their phase difference to increase the utility (respectively increase the phase difference when u_ij_(t)<0). To compute the cumulative synergy of an interaction, since s_i_(t) and s_j_(t) vary over time, we need to integrate utility with respect to time, over the course of a cycle:

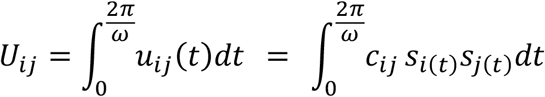

However, s_i_(t) is very difficult to integrate, because it depends on *I*_*i*(*t*)_, which itself depends on multiple other functions (s_k_(t), s_l_(t), s_m_(t), etc.). A solution can come from the fact that*I*_*i*(*t*)_ only has a short transient effect on s_i_(t) over the course of a cycle. The dynamics of s_i_(t) is nearly sinusoidal, and its magnitude can be approximated by γ_*i*(*t*)_, varying between 0 and +1/-1 over a cycle. Thus, utility can be approximated as:

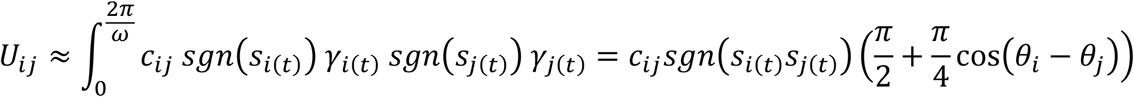

by considering that sgn(s_i_(t)) is constant over the course of a cycle. Hence the following question: how would the utility of an interaction change when the phase difference changes? From the previous approximation, we can compute:

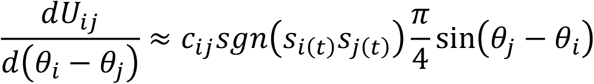

If we consider that a unit *i* tries to increase the synergy of the interactions with positive utility and decrease the synergy of the interactions with negative utility, *θ*_*i*_ values should follow the following equation of motion:

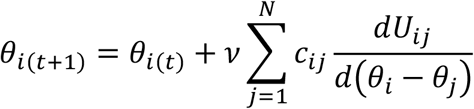

with ν being the constant controlling the rate of change of phase. By introducing the approximation of U_ij_ defined above, this equation of motion becomes Eq. 5.

## Footnotes

1 The utility of a unit is just the weighted sum of constraints acting on it (Watson, Mill, Buckley 2011a, 2011b) i.e.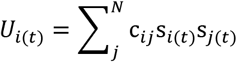. The energy function, Eq. 2, is equivalent to the negation of total utility or the sum of individual utilities.

2 Biological organisms often exhibit many levels of agency but here we focus on a simple case of just one level of emergent agency to understand and clarify sufficient dynamics. Also natural biological interactions will most likely have interactions of various different strengths, exhibiting complex (sometimes unresolvable) frustrations – whereas, here we are using a problem with neat, non-overlapping modules (‘countries’), and *consistent* constraints (i.e. there exists a configuration where all constraints can be resolved simultaneously). Although these modules clearly prescribe the boundaries of the collectives that form, it is still necessary for a bottom-up process to identify where they are and form relationships, with the necessary members, that facilitate higher-level agency.

3 This is used in many models of development, including cell-cell communication (Ferrell 2012, Fooladi, Moradi et al. 2019, Matsushita & Kaneko 2020, Thompson 2021).

